# Microbiome engineers: Grazers, browsers, and the phytobiome stampede

**DOI:** 10.1101/087494

**Authors:** K. A. Garrett, E. E. Frank, S. P. Dendy, J. F. Leslie, A. A. Saleh

## Abstract

We define *microbiome engineers* as species that modify the microbiome associated with other host species via changes in the physical environment, potentially including the creation of dispersal networks for microbiome consortia across multiple hosts. Grazers such as bison are indirect plant microbiome engineers through alteration of the structure of plant communities in grasslands and forests. They also can directly engineer plant microbiomes if they distribute microbial consortia. Direct engineering may include simpler examples such as the role of truffle-eating animals in structuring forest mycorrhizal communities, as well as more complex roles in structuring the evolution of bacterial, fungal, and viral microbiome networks. Grazers and browsers may have important historic and current roles in engineering microbiomes by (a) stabilizing and homogenizing microbiomes in their preferred plant species, and (b) selecting for microbiomes in their preferred plant species that are differentiated from microbiomes in plant species they rarely consume.

---

Ecosystem engineers are species that manipulate the physical environment, apart from trophic interactions, in ways that may affect numerous other species (Jones *et al.*, 1997). Beavers are a classic example of an ecosystem engineer, constructing dams that support an ecological community dependent on the resulting wetlands (Wright *et al.*, 2002) and connectivity among wetlands (Hood and Larson, 2015). An individual plant may be considered an ecosystem engineer through effects on the environment, including production of root exudates (Peiffer *et al.*, 2013). We define a *microbiome engineer* as a species that directly and/or indirectly modifies the spatial and temporal structure of the microbiome associated with other host species through changes in the physical environment, potentially including creation of networks for dispersal of microbial consortia across multiple host individuals (Fig. 1). This definition includes effects that remove limitations to dispersal, in addition to effects on availability of resources (such as nutrients) emphasized in definitions of ecosystem engineers. In contrast to more specific vectors, a microbiome engineer influences larger microbial consortia. Humans may be, or may become, the most influential microbiome engineer, developing synthetic communities and even synthetic life forms (Mueller and Sachs, 2015).

**Figure 1.**
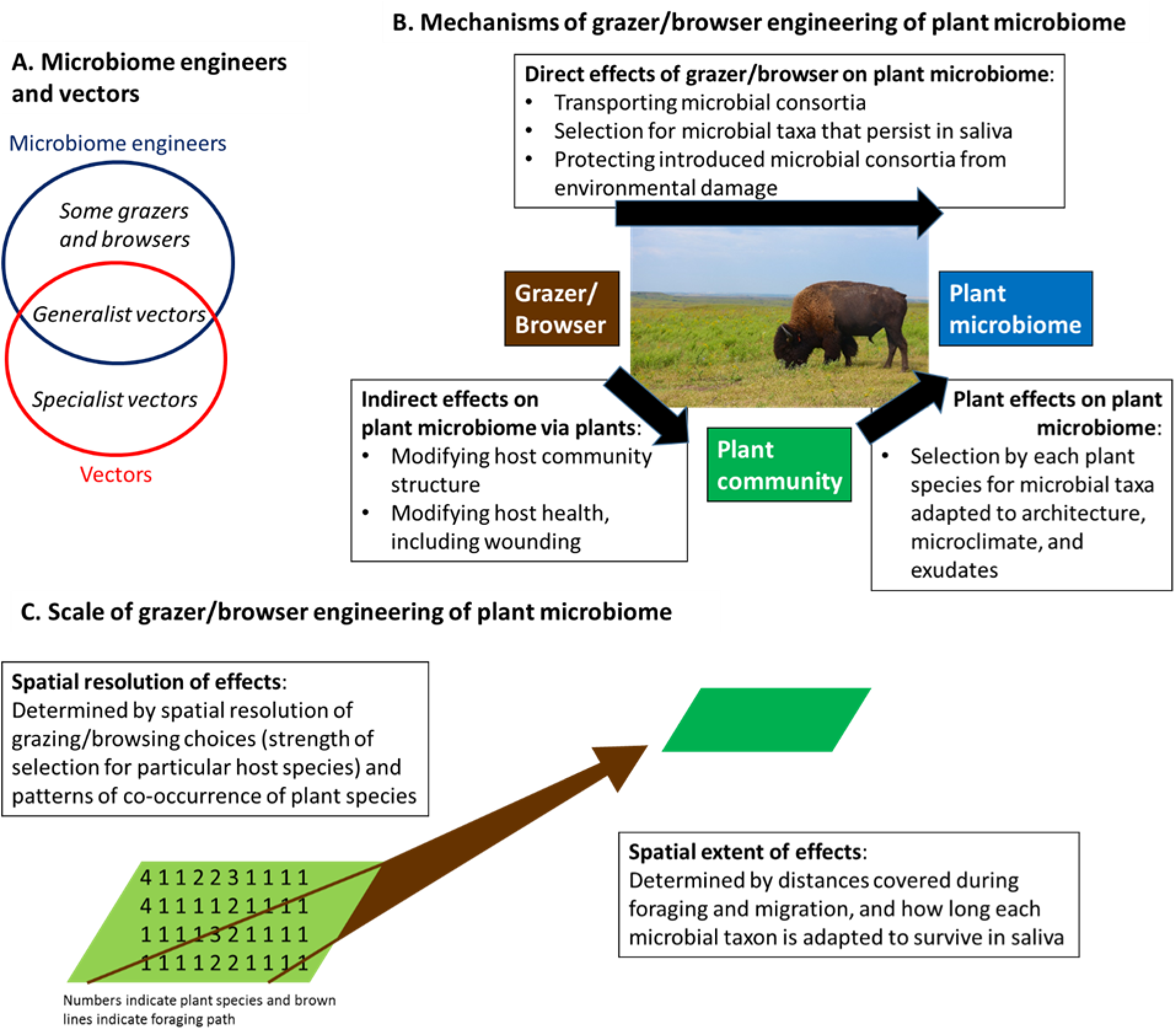
A. Microbiome engineers as defined here affect microbial consortia, rather than only individual microbial taxa (although a specialist vector could have indirect effects on the phytobiome by modifying host health). B. Grazers and browsers are particularly well-positioned to be microbiome engineers for living plant populations and communities because they create an entry point to plants without killing plants. Many grazed/browsed plants are well-adapted to recover and may regrow with the modified phytobiome. The mechanisms of grazer/browser effects on plant microbiomes may be direct or indirect, via effects on plants. C. Grazers/browsers may engineer plant microbiomes across individuals within a plant species, and across plant species. (Photo of bison by B. Van Slyke)

The goal of this paper is to develop the concept of the microbiome engineer, particularly in the context of vertebrate plant grazers and browsers, to illustrate the key role such animals may play, and how high-throughput sequencing provides a new platform for understanding engineering of whole consortia rather than simply a few taxa with known ecosystem roles.

Some links between mammals and plant-associated microbes are well-documented (Johnson, 1996), such as how flying squirrels consume and spread ectomycorrhizal fungi and nitrogen-fixing bacteria, structuring forest communities (e.g., (Li *et al.*, 1986; Lehmkuhl *et al.*, 2004)). Invasive mammals may facilitate the invasion of fungi and trees (Wood *et al.*, 2015).

Grazers and browsers may likewise spread many microbes through feces (Supplemental Materials), but the process of grazing and browsing has particular potential for direct microbiome engineering because it can introduce microbial consortia directly to wounded plant organs that are adapted to recover. Transfer through grazing and browsing may also provide protection against desiccation via saliva, and nutrients, for the microbial consortia introduced into plant wounds. Microbes moved by microbiome engineers are co-evolved consortia, so there may be a high probability of successful establishment. High-throughput sequencing opens the door to studies of the movement of large assemblages of microbial taxa.

The American bison is well-known for its direct effects on plant communities (Hartnett *et al.*, 1996; Knapp *et al.*, 1999; Towne *et al.*, 2005; Borer *et al.*, 2009), thus must have corresponding important *indirect* effects on plant microbiomes, and may have *direct* effects by moving microbes when grazing (Fig. 1). Cow saliva can disperse the fungus *Ascochyta paspali* in *Paspalum dilatatum*, causing disease (Williams and Price, 1989). Bison have the potential to act as direct microbiome engineers of prairies, because they maintain multiple microbial species in their saliva, and thus may redistribute these microbial consortia through grazing (new data in Supplemental Material). Grazing and browsing species may act as a hub or bridge node in networks of microbial taxon association, where key microbial consortia associated with these animals have a cascade of effects on other taxa (Faust and Raes, 2012; Poudel *et al.*, 2016).

Some functions of the taxa we recovered from bison saliva are known. *Fusarium* spp. can produce mycotoxins, where the effects of mycotoxins in natural systems are mostly unknown. In general, monogastric animals are much more sensitive to mycotoxins than are ruminants such as bison. Species recovered from bison saliva play a role in regulating plant growth (Harman *et al.*, 2004), and endophytic strains of *Fusarium* capable of synthesizing gibberellic acid have previously been recovered from tallgrass prairie (Leslie *et al.*, 2004). An isolate of the plant pathogen *Macrophomina phaseolina* from bison saliva was genetically similar to an isolate from tallgrass prairie plants (Saleh *et al.*, 2010). New roles for microbial species will continue to be discovered, *e.g*., heat tolerance conferred by fungi infected with a virus (Márquez *et al.*, 2007). Our data (Supplemental Material) focused on fungi, but grazers such as bison also could disperse plant-associated viruses, bacteria, and nematodes.

Grazers and browsers may be important agents in shaping the spatial and temporal distribution of plant-associated microbes. The combination of foraging decisions by grazers such as bison (Courant and Fortin, 2010), and the pattern of grassland plant species co-occurrence (Cox *et al.*, 2013), will help to determine the likelihood that particular plant species are grazed in close enough temporal proximity to help homogenize their microbiomes. The importance of microbiome engineers is determined by a number of such factors (Fig. 2), including whether dispersal of microbes would otherwise be a limiting factor for community assembly (Martiny *et al.*, 2006). If microbes dispersed by grazers and browsers produce secondary metabolites in plants that alter the palatability of a plant, this feedback may produce patterns that are self-reinforcing. Differences in grazing patterns and oral microbiome selection by wild and domestic grazers and browsers could produce complex, overlapping microbiome engineering effects.

**Figure 2.**
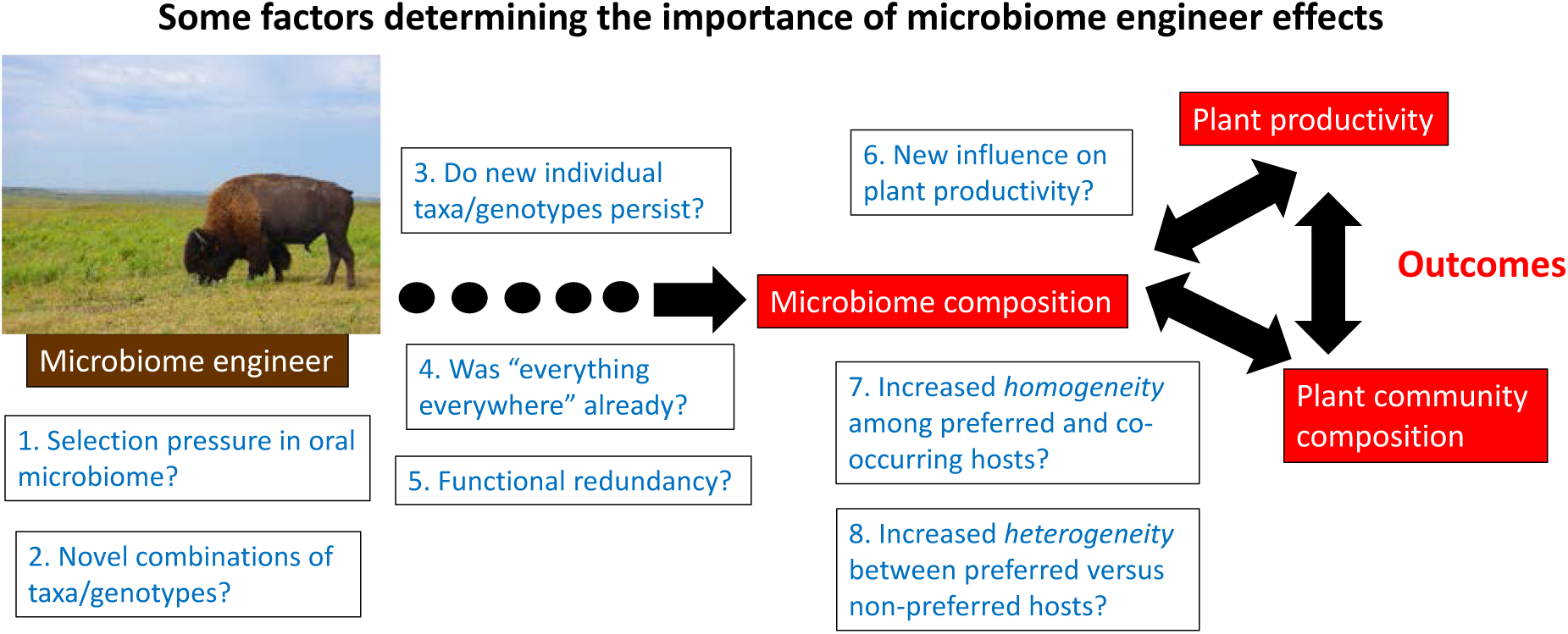
Potential outcomes of direct microbiome engineering, and scenarios producing more or less important effects

1. Does selection pressure in the oral microbiome, e.g., due to abundance and properties of saliva, change the microbial consortium?
2. Does dispersal of microbial consortia produce new communities of taxa/genotypes that would not assemble in plant microbiomes without grazing/browsing?
3. Do new individual taxa/genotypes introduced to the phytobiome persist?
4. Is it that “everything is everywhere” already, or does dispersal of microbial consortia (especially into plant wounds) enable plant microbiome assembly?
5. Are new taxa/genotypes introduced to the plant microbiome functionally redundant with pre-existing microbiome?
6. Do new taxa/genotypes or new combinations of taxa/genotypes influence plant productivity?
7. Does engineering result in increased microbiome homogeneity among individuals in preferred plant species, and among plant species that tend to co-occur with them?
8. Does engineering result in increased microbiome heterogeneity among preferred versus non preferred plant species? (Photo of bison by B. Van Slyke)

A wide range of global vertebrate and invertebrate herbivores may be direct plant microbiome engineers, including animals such as rabbits, giraffes, marsupials, geese, and possibly even aquatic grazers such as sea turtles and manatees. One key feature of ecosystem engineers is that their direct effects can outlive an individual engineer (Hastings *et al.*, 2007). Microbiome engineers may influence the evolutionary trajectories of plant microbiomes long after engineers have migrated to other areas (Odling-Smee *et al.*, 2013), and may re-introduce taxa that are periodically or patchily less-favored, a process particularly important in annual grasslands. The ability of bison saliva to maintain viable plant-associated microbes suggests a potential role in long-distance dispersal and phytobiome engineering for other migratory herbivores, *e.g*., wildebeest and Cape buffalo in Africa, caribou and musk ox in the Arctic, and kangaroos in the Australian Outback. Large mammal migrations may have engineered the geographic structure of the plant microbiome across many of the world’s grasslands, savannahs, and forests.

## Acknowledgements

KAG developed the microbiome engineer concept with input from all authors; KAG and JFL conceived the bison saliva microbiome experiment; AAS, EEF, and SPD implemented and analyzed the bison saliva microbiome experiment; KAG and AAS wrote the manuscript with input from all authors. We appreciate helpful discussions with P. Garfinkel, A. Joern, R. Poudel, T. Todd, and E. Towne, and support by the NSF under Grants DEB-0130692 and DEB-0516046, by NSF grant EF-0525712 as part of the joint NSF-NIH Ecology of Infectious Disease program, by the NSF Long Term Ecological Research Program at Konza Prairie, by the Kansas Agricultural Experiment Station, and by the University of Florida.

## Supplemental Material: Bison oral microbiome

### Background Information: Bison and Tallgrass Prairie

Bison are known to have direct effects on the structure of tallgrass prairie plant communities (Hartnett *et al.*, 1996; Knapp *et al.*, 1999; Towne *et al.*, 2005). In studies at Konza Prairie Biological Station (KPBS, Manhattan, Kansas, USA), bison select grazing areas based on four characteristics: a high population of the dominant tallgrass prairie plant species, lower forb richness and diversity, lower plant species diversity, and high grass to forb ratios (Vinton, 1993). Within bison grazing lawns, bison prefer particular plant species and areas in the lawns with regrowth in grazed patches of high quality (Weaver and Tomanek, 1951; McNaughton, 1984). The composition of bison diets changes as availability changes with the seasons. In early spring and fall they spend more time in areas that are dominated by C_3_ plants, while in the summer most of their time is spent in areas dominated by C_4_ grasses (Vinton *et al.*, 1993). Grazers may indirectly support the spread of plant pathogens if they increase the abundance of highly-competent hosts (Borer *et al.*, 2009). If they also spread plant-associated microbes, their grazing patterns may help to structure the geographic patterns of plant microbiomes across time.

Plant-associated microbes are dispersed by numerous taxa, with the most widely documented vectors being arthropods. Arthropods vector pathogens including bacteria, viruses, fungi, and phytoplasmas. Other vectors include fungi, nematodes, and earthworms, while vertebrates are less well-known as vectors of plant-associated microbes. Cow saliva can disperse the fungus *Ascochyta paspali*, with inoculation of healthy plants by saliva containing fungal conidia resulting in leaf blight disease in *Paspalum dilatatum* plants (Williams and Price, 1989). Saliva effects on plants also have been evaluated in sheep, where saliva had positive effects on plant growth (Teng *et al.*, 2010). Native grazers such as bison may play a major and underappreciated role in the distribution of plant-associated microbes in grasslands.

The effects of bison grazing on plants have been evaluated in terms of the chemical composition of bison saliva, and whether their saliva contains plant growth regulators (Detling *et al.*, 1980; Detling *et al.*, 1981). Compensatory plant growth may occur in response to herbivory (McNaughton, 1983). Some of the effects of saliva observed in the past may be due to microbes contained in the saliva. Microbes in the saliva of grazers could be effectively inoculated into a plant through grazing, as some saliva would remain on the plant wounds created by grazing and might protect microbes from desiccation, and might also supply some nutrients. Microbes within the saliva would be well placed to enter through the wounds and bypass many of the barriers found in intact plant leaves and stems.

Plant-associated microbes play multiple roles in tallgrass prairies. Mycorrhizal associations are important for the success of the dominant grass species (Hetrick *et al.*, 1988; Hartnett and Wilson, 1999), and other fungi may positively influence plant growth (Harman *et al.*, 2004). The role of plant pathogens in tallgrass prairie is less understood. Many pathogens occur in tallgrass prairie (Tiffany *et al.*, 1990) with their prevalence both influenced by and influencing plant community composition (Mitchell *et al.*, 2002; Barnes *et al.*, 2005). Fungicide application can increase tallgrass prairie grass productivity and yields, suggesting the potential importance of pathogens in limiting grass productivity (Mitchell, 2003; Dickson and Mitchell, 2010). New grassland fungus species, *e.g*., *Fusarium konzum* (Zeller *et al.*, 2003), are likely to be described as more attention is given to microbes in natural systems.

We sampled bison saliva for viable fungal propagules, to demonstrate the potential for bison to spread microbial consortia while grazing.

### Methods in Bison Saliva Analysis

*Sampling*. A herd of bison maintained at Konza Prairie Biological Station (KPBS) (Towne, 1999) grazes native tallgrass prairie. Each autumn the bison are assembled for weighing, offering an opportunity to sample their saliva. While briefly confined in the corral during roundup, the bison had access to dried native prairie hay from KPBS that had been baled in the summer. Thus, it is possible that the animals’ access to the native prairie hay biased our sample toward fungal species that can survive in dried prairie plant material. (During winter, bison naturally feed on plant material dried in the field.) We sampled saliva from 51 bison (33 females and 18 males) with a single sterile veterinary swab per animal as individual animals were held in a chute.

*Isolation of Viable Fungi*. Cotton swabs were streaked onto peptone/PCNB medium (Leslie and Summerell, 2006). This medium is semi-selective for *Fusarium* spp. and other fungi that can tolerate PCNB, streptomycin, and neomycin, and utilize an amino acid mixture as the sole source of carbon and nitrogen. Swabs were then rotated 180° and streaked onto *Macrophomina* selective media (Mihail, 1992). Plates were incubated at room temperature (25±2°C) for 7-10 days. The extra step of culturing before sequencing makes it possible to say definitively that the fungi were viable, whereas direct sequencing of DNA from the saliva also would detect DNA from dead microbes.

Strains of interest were mass-transferred to Potato-Dextrose agar (PDA, Difco, Detroit, MI, USA) slants and stored at 4°C for further purification and identification. For purification, strains were transferred from PDA slants to Petri dishes containing peptone/PCNB medium or PDA plates. Plates were incubated at 22°C with a 12 hr:12 hr light:dark cycle for up to two weeks. Spores from colonies growing on the peptone/PCNB medium were streaked on a water agar slab for microscopic examination and for subculturing single conidia to ensure culture purity. Three to four spores were separated from each streak by micromanipulation. Slabs were incubated overnight at 25°C. Germinated single spores were cut out of the agar slab under dissecting microscope (50×) and transferred to Petri dishes containing Czapek’s complete medium (Leslie and Summerell, 2006). Cultures were incubated at 22°C with a 12 hr:12 hr light:dark cycle.

For morphological identification, pure colonies were subcultured to appropriate media and fungal strains were identified according to Leslie and Summerell (2006) and Barnett and Hunter (1998). DNA was extracted from cultures that had been grown on liquid complete medium and incubated on an orbital shaker (150 rpm) at room temperature for 48-72 hours by using a cetyltrimethylammonium bromide (CTAB) procedure (Leslie and Summerell, 2006).

*Sequencing*. Two genes were amplified and sequenced to help identify strains to species level: β-tubulin (*tub-2*) and the internal transcribed spacer (ITS1 and ITS2) region of nuclear rDNA gene (ITS-rDNA) (White *et al.*, 1990; O’Donnell *et al.*, 2000; Saleh *et al.*, 2010). The DNA sequences were edited and aligned with BioEdit software (http://www.mbio.ncsu.edu/Bioedit/bioedit.html). Cleaned DNA sequences were searched against the GenBank database to identify identical or closely related fungi to assign strains to a genus and species wherever possible.

### Results and Discussion of Bison Saliva Analysis

A range of fungal taxa were recovered, with 30 fungal cultures identified to species level (Table S1). Among the dematiaceous fungi recovered from bison saliva, one isolate was identified as *Macrophomina phaseolina* and had the same haplotype as the MPKS327 strain previously recovered from *Helianthus* sp. growing in a Kansas tallgrass prairie (Saleh *et al.*, 2010). The fungal species most frequently recovered (15/65 isolates) was *Phoma sorghina*, which can cause foliar diseases in grasses and other plants. Among the hyaline-hyphae fungi, isolates from three *Fusarium* species complexes were recovered: *Fusarium equiseti, Fusarium oxysporum*, and *Fusarium solani*. Members of these *Fusarium* species complexes are all ubiquitous soil inhabitants. Their presence may be due to an endophytic or pathogenic plant association, or they may be non-pathogens present temporarily or growing as saprophytes on plant surfaces. While some of the recovered species are well-known plant pathogens, such as *M. phaseolina, Stagonosporopsis cucurbitacearum* and *P. sorghina*, others may be pathogens, symbionts or hitchhikers, with their role dependent on the plant host and environmental conditions (Newton *et al.*, 2010).

Viable fungi were recovered from 41/51 bison on peptone/PCNB medium, and from 14/51 bison on *Macrophomina* selective medium. There were 65 cultures for evaluation following the single-spore subculturing process. Based on morphological characterization, the recovered fungal strains were divided into two groups: fungi with hyaline hyphae and fungi with dark-colored hyphae (dematiaceous fungi). We amplified and sequenced both the *tub-2* and ITS-rDNA regions from 43/65 fungal strains. For the remaining 22 strains either the ITS-rDNA (17 strains) or *tub-2* (5 strains) region was amplified and sequenced. When these DNA sequences were searched against the GenBank database, 57/65 fungal cultures were identified to the genus level with an average 99% identity for ITS-rDNA regions and 90% identity for *tub-2* regions. In addition, 30 out of the 57 fungal cultures were identified to the species level (Tables S1).

Among the hyaline-hyphae fungi, members of three Fusarium species complexes were recovered: *F. equiseti* (seven strains with ITS-rDNA and tub-2 sequences 100% and 95% identical to sequences already deposited in GenBank, respectively), *F. oxysporum* (four strains with sequences 100% identical to those deposited in GenBank for both sequenced regions), and *F. solani* (one strain with sequences 99% identical to those deposited in GenBank for both regions). Fungal strains identified as *Plectosphaerella* sp. (5 strains), *Verticillium* sp. (4 strains), *Aspergillus tubingensis* (1 strain), *Acremonium* sp. (1 strain), and *Colletotrichum* sp. (1 strain) also were recovered. While identification to genus is not sufficient to evaluate the ecological role for these taxa, all of them could be plant pathogens. There were eight strains with less than 90% DNA sequence similarity to any of the DNA sequences available in GenBank.

Recovery of multiple species of viable fungi indicates the potential for bison to directly engineer grassland microbiomes, in addition to their indirect effects on the microbiome through structuring plant communities. The common impression that soilborne and endophytic fungi are limited to very gradual dispersal, through soil movement and plant growth, must be modified to take into account dispersal by bison and probably other grazers. Bison have historically been one of the most abundant large land mammals known, with perhaps 50 million bison ranging from Canada to Mexico around 2000 years ago. Bison travel approximately 3 km a day in typical grazing patterns, but have historically moved much greater distances in seasonal migrations (Meagher, 1986). The range of the ~ 10,000 free-ranging animals today (Meagher, 1986; Nowak, 1999; Lott, 2002) is much more limited and does not include the vast majority of the Great Plains region where they roamed freely until 100-150 years ago, dispersing microbes as they grazed.

Bison have the potential to directly engineer both plant microbiomes and soil microbiomes more broadly. Soilborne microbes could be moved over the period of several days or longer, including movement in bison fur (from dust wallows) and hooves, and carried a considerable distance, especially to other wallows. Plant-associated microbes could be moved in saliva across distances covered in one to a few days, or potentially longer, while they remained viable in an animal’s mouth. If the microbes were swallowed, those that survive the animal’s digestive system could be dispersed in the dung. Members of all three of the *Fusarium* species complexes recovered, for example, produce thick-walled chlamydospores that could transit the bison’s digestive tract. 96% of the DNA sequences identified from the dung of pronghorn antelope (*Antilocapra americana*), which also graze the Great Plains, were categorized as *Phoma* species (Herrera *et al.*, 2011). The predominance of viable *Phoma* propagules among the fungi we recovered from bison saliva could indicate that this important group of plant pathogens has evolved in concert with large herbivores in the Great Plains, helping to insure dispersal throughout the region. The profile of bacteria observed in bison dung suggests differences that may reflect bison health status (Weese *et al.*, 2014), and bison health status might help determine which microbes survive in saliva or dung.

Unlike microbe transport in soil or deposited in dung, bison saliva is ideally positioned for plant infection. As the animals graze, they create wounds to which saliva would be simultaneously applied. Organisms transmitted in saliva thus need fewer long-term survival structures and lesser abilities to abrogate or bypass plant defenses because they would effectively be injected directly into the plant vascular system. These microorganisms could have adapted to short-term and possibly long-term survival in the bison oral microbiome in much the same way that other microorganisms have adapted to survive in much better understood arthropod vectors.

Global grassland microbiomes likely retain some spatial structure based on historic grazing patterns, while more recent replacements of wild grazers with domesticate animals may continue to modify these patterns.

---

**Table S1.** Viable fungi recovered from bison saliva, identified using morphological and molecular markers

| Number of<br>isolates | Fungal taxon |
| --- | --- |
| 15 | *Phoma sorghina* |
| 4 | *Fusarium oxysporum* |
| 2 | *Stagonosporopsis cucurbitacearum* |
| 1 | *Aspergillus tubingensis* |
| 1 | *Fusarium solani* |
| 1 | *Macrophomina phaseolina* |
| 15 | *Phoma* species complex |
| 7 | *Fusarium equiseti* species complex |
| 5 | *Plectosphaerella* sp. |
| 4 | *Verticillium* sp. |
| 1 | *Acremonium* sp. |
| 1 | *Colletotrichum* sp. |
| 8 | Ascomycete |

